# Apropos of Universal Epitope Discovery for COVID-19 Vaccines: A Framework for Targeted Phage Display-Based Delivery and Integration of New Evaluation Tools

**DOI:** 10.1101/2021.08.30.458222

**Authors:** Christopher Markosian, Daniela I. Staquicini, Prashant Dogra, Esteban Dodero-Rojas, Fenny H. F. Tang, Tracey L. Smith, Vinícius G. Contessoto, Steven K. Libutti, Zhihui Wang, Vittorio Cristini, Paul C. Whitford, Stephen K. Burley, José N. Onuchic, Renata Pasqualini, Wadih Arap

**Affiliations:** Rutgers Cancer Institute of New Jersey, Newark, NJ 07101; Division of Cancer Biology, Department of Radiation Oncology, Rutgers New Jersey Medical School, Newark, NJ 07103; Mathematics in Medicine Program, Houston Methodist Research Institute, Houston, TX 77030; Department of Physiology and Biophysics, Weill Cornell Medical College, New York, NY 10022; Center for Theoretical Biological Physics, Rice University, Houston, TX 77005; Department of Physics, Institute of Biosciences, Humanities and Exact Sciences, São Paulo State University, São José do Rio Preto, SP 15054, Brazil; Rutgers Cancer Institute of New Jersey, New Brunswick, NJ 08901; Department of Surgery, Rutgers Robert Wood Johnson Medical School, New Brunswick, NJ 08901; Department of Imaging Physics, University of Texas M.D. Anderson Cancer Center, Houston, TX 77030; Physiology, Biophysics, and Systems Biology Program, Graduate School of Medical Sciences, Weill Cornell Medicine, New York, NY 10022; Department of Physics and Center for Theoretical Biological Physics, Northeastern University, Boston, MA 02115; Department of Chemistry and Chemical Biology, Rutgers, The State University of New Jersey, Piscataway, NJ 08854; RCSB Protein Data Bank and Institute for Quantitative Biomedicine, Rutgers, The State University of New Jersey, Piscataway, NJ 08854; RCSB Protein Data Bank, San Diego Supercomputer Center and Skaggs School of Pharmacy and Pharmaceutical Sciences, University of California San Diego, La Jolla, CA 92067; Department of Biosciences, Rice University, Houston, TX 77005; Department of Chemistry, Rice University, Houston, TX 77005; Department of Physics and Astronomy, Rice University, Houston, TX 77005; Division of Hematology/Oncology, Department of Medicine, Rutgers New Jersey Medical School, Newark, NJ 07103

**Author notes:** These authors jointly supervised this work (Renata Pasqualini, Wadih Arap). Correspondence should be addressed to Wadih Arap or Renata Pasqualini, Rutgers Cancer Institute of New Jersey, Newark, NJ 07101.,.

## Abstract

Targeted bacteriophage (phage) particles are potentially attractive yet inexpensive platforms for immunization. Herein, we describe targeted phage capsid display of an immunogenically relevant epitope of the SARS-CoV-2 Spike protein that is empirically conserved, likely due to the high mutational cost among all variants identified to date. This observation may herald an approach to developing vaccine candidates for broad-spectrum, towards universal, protection against multiple emergent variants of coronavirus that cause COVID-19.

## Main

The ongoing Coronavirus Disease 2019 (COVID-19) global pandemic continues to pose an unprecedented health threat to humankind and the potential for evolution of more infectious and/or virulent SARS-CoV-2 variants with resistance to vaccines remains a considerable concern.^1^ Within ∼12 months after SARS-CoV-2 first emerged clinically, the United States Food & Drug Administration (FDA) began issuing Emergency Use Authorization (EUA) for COVID-19 vaccines for the general population. Less than nine months out from their initial deployment, we have been experiencing two setbacks in the fight against the virus. First, the Delta “Variant of Concern” (VOC) is proving to be more transmissible and more virulent than the original Alpha VOC that swept around the globe in late 2020. Second, immunological protection unfortunately appears to wane in the months following either initial infection or full vaccination. Though the FDA just granted full approval for the Pfizer-BioNTech vaccine,^2^ COVID-19 vaccines in general have been reported to have decreased effectiveness against Delta variant infection and symptomatic illness.^3^

SARS-CoV-2 has a linear positive-sense RNA genome that encodes four main structural proteins.^4^ The Spike (S) protein, which is the main target of neutralizing antibodies generated following infection by SARS-CoV-2,^5^ has formed the basis of nearly all first-generation COVID-19 vaccines.^6,7^ A recent massive analysis of more than 300,000 viral genomes has suggested that mutations within the S protein represent one of the main pathways of adaptive evolution in SARS-CoV-2,^8^ thereby raising concerns about potential for resistance to neutralizing antibodies acquired through either vaccination or previous COVID-19 infection.^9-11^ Recent developments have supported the concept of “inescapable” monoclonal antibodies with broad activity against multiple variants of SARS-CoV-2^12^; however the ongoing genetic evolution of the virus raises uncertainty regarding the possibility of new viral mutations that allow immune escape. Moreover, given that some proportion of the viral mutations may also alter infectivity, disease severity, interaction with host immunity, and resistance to antiviral drugs,^13^ both the World Health Organization (WHO)^14^ and the United States Centers for Disease Control & Prevention (CDC)^15^ have actively been monitoring at least 21 new variants of SARS-CoV-2. By definition, the ones with greater numbers of mutations are associated with high transmissibility and thus labeled VOCs, while those with altered biological features or immunogenicity are termed “Variants of Interest” (VOIs).^9,13^ In the face of the highly transmissible Delta VOC that has recently caused a rapid rise in infection cases and concerns related to the efficacy of current vaccines, multiple countries are now recommending yet an additional dose (“booster”) of vaccine for fully vaccinated but immunocompromised individuals^16^ and for those with normal immune systems more than eight months post vaccination.

Rapid evolution of the SARS-CoV-2 genome^13^ highlights the need for developing a single vaccine (or a small number of vaccines) with a broad, ideally universal, spectrum of activity against viral variants. To achieve these desirable attributes, development candidate vaccine antigens should be: (i) immunogenic, (ii) highly conserved (*i*.*e*., free of non-conserved missense mutations and/or single-residue deletions), (iii) capable of recapitulating its endogenous conformation when displayed in heterologous contexts, and (iv) surface solvent-exposed and thus suitable for recognition by antibodies and other ligands from either the humoral or cellular immune response. Several recent studies with different strategies such as artificial intelligence/machine learning (AI/ML),^17^ phage display-based immunoprecipitation and sequencing (PhIP-Seq) technology,^18^ or systematic site-directed mutagenesis^19^ scanned and predicted mutational hotspot regions for design of broad-spectrum vaccines, diagnostics, and antibody-based therapies. Notwithstanding insights into epitope mapping, it remains unclear whether these polypeptide chain segments are evolutionary stable and, therefore, suitable to control the spread of emerging COVID-19 variants worldwide.

In previous work, we have established alternative vaccine strategies based on targeted pulmonary immunization with ligand-directed phage constructs.^20^ Phage particles are generally harmless to humans, but induce a potent non-specific humoral response; they are inexpensively produced and distributed at industrial scale with no temperature-controlled supply cold-chain requirement,^21^ which represent major impediments to moving first-generation COVID-19 vaccines into resource-limited settings. To drive the phage-based vaccine concept towards clinical translation, we have demonstrated the concept of an aerosol-delivered phage-based vaccine prototype.^20,21^ Phage particles were engineered to display a highly-immunogenic cyclic decapeptide epitope of the SARS-CoV-2 S protein (residues C662–C671, sequence CDIPIGAGIC) (hereafter referred to as the C662–C671 epitope) fused with a lung-directed motif that facilitates uptake and distribution after targeted aerosol delivery.^21^ We showed that the C662– C671 epitope closely recapitulates its native conformation within the full-length S protein when exposed on the surface of phage and elicits a strong and specific antibody response in mice. This experimental evidence strongly suggests that such a strategy fulfills the functional attributes for induction of a broad immune response to existing and/or new variants.^21^

Herein, we investigated the C662–C671 epitope further by serially performing comparative sequence, structural, simulation, and computational analyses of the recently emerging mutational impact of VOCs and VOIs in the context of our candidate phage-based vaccine prototype.^20,21^ Our analyses reveal that the C662–C671 epitope indeed achieves the aforementioned structural and functional requirements and, therefore, might represent a unique opportunity for the development of a vaccine that would elicit robust and durable protection against known and emergent SARS-CoV-2 variants.

To examine the nature of mutational selective pressure on SARS-CoV-2, we analyzed S protein sequences from all four VOCs and four VOIs (as deemed by the CDC, accessed on August 30, 2021).^15^ We observed that although these eight variants harbor mutations at 48 amino acid positions distributed throughout the 1,273-residue protein (**Table 1**), the C662–C671 epitope is strictly conserved, an attribute that could be exploited towards the development of a broad-spectrum, possibly universal, COVID-19 vaccine. Of note, mutations in the VOCs (Alpha, Beta, Gamma, and Delta) are primarily clustered in the N-terminal domain (NTD) and the receptor-binding domain (RBD) of S protein (**Fig. 1a**). Based on the antibody repertoires identified in plasma of SARS-CoV-2-infected individuals, both RBD and NTD are the targets of various neutralizing antibodies, with RBD being immunodominant.^13^ Moreover, selection pressure on the virus from neutralizing antibodies generated *via* natural infection, vaccines, or antibody-based therapies is also consistent with the rapid rise in mutations within the RBD, which is highly tolerant of immune-evading protein changes based on evolutionary modeling.^22^ The C662–C671 epitope, which is located outside the NTD and the RBD, has also been shown to be targeted by neutralizing antibodies.^23-25^ Our analysis, however, shows that this epitope does not appear to be mutation-prone. While studies with patient convalescent serum samples suggest that antigen escape from neutralizing antibodies likely drives the rise and dominance of new variants,^13^ conserved epitopes across different strains, such as C662–C671, may help guide development of a vaccine candidate against different SARS-CoV-2 variants. Notably, none of the additional variants monitored by the WHO (as accessed on August 30, 2021), including the Lambda VOI together with variants not as yet assigned a Greek alphabet letter such as B.1.427, B.1.429, R.1, B.1.466.2, B.1.621, B.1.1.318, B.1.1.519, C.36.3, B.1.214.2, B.1.1.523, B.1.619, and B.1.620,^14^ harbor missense mutations and/or amino acid deletions affecting the C662–C671 epitope. Moreover, such mutations within C662– C671 have only been identified in merely 1,313 instances of at least 2,659,222 sequenced (a total of ∼0.05% for the 10-residue epitope) SARS-CoV-2 isolates worldwide.^26^ Collectively, these empirical results suggest that C662–C671 epitope might represent a privileged motif for vaccine development and perhaps even a potential target for inescapable antibodies.^12^

**Table 1:** List of amino acid substitutions or deletions for SARS-CoV-2 VOCs and VOIs

| Variants of Concern (VOCs) |  |  |  | Variants of Interest (VOIs) |  |  |  | Top S protein mutation sites |  |
| --- | --- | --- | --- | --- | --- | --- | --- | --- | --- |
| Alpha | Beta | Gamma | Delta | Eta | Iota | Kappa | B.1.617.3 | Residue site | # variants |
| H69Δ | D80A | L18F | T19R | A67V | L5F | (T95I) | T19R | 614 | 8 |
| V70Δ | D215G | T20N | (V70F) | H69Δ | (D80G) | G142D | G142D |  |  |
| Y144Δ | L241Δ | P26S | T95I | V70Δ | T95I | E154K | L452R | 484 | 7 |
| (E484K) | L242Δ | D138Y | G142D | Y144Δ | (Y144Δ) | L452R | E484Q |  |  |
| (S494P) | A243Δ | R190S | E156Δ | E484K | (F157S) | E484Q | D614G | 452 | 4 |
| N501Y | K417N | K417T | F157Δ | D614G | D253G | D614G | P681R |  |  |
| A570D | E484K | E484K | R158G | Q677H | (L452R) | P681R | D950N | 681 | 4 |
| D614G | N501Y | N501Y | (A222V) | F888L | (S477N) | Q1071H |  |  |  |
| P681H | D614G | D614G | (W258L) |  | E484K |  |  | 70, 95, | 3 |
| T716I | A701V | H655Y | (K417N) |  | D614G |  |  | 142, 144, |  |
| S982A |  | T1027I | L452R |  | A701V |  |  | 417, 501, |  |
| D1118H |  |  | T478K |  | (T859N) |  |  | 950 |  |
| (K1191N) |  |  | D614G |  | (D950H) |  |  | 19, 69, 80, | 2 |
|  |  |  | P681R |  | (Q957R) |  |  | 157, 701 |  |
|  |  |  | D950N |  |  |  |  |  |  |

Notes: Variants were deemed as VOCs or VOIs by the CDC (as accessed online on 08/30/2021).<sup>15</sup> Mutations listed in parentheses are present in only some strains of the corresponding variant.

**Fig. 1.**
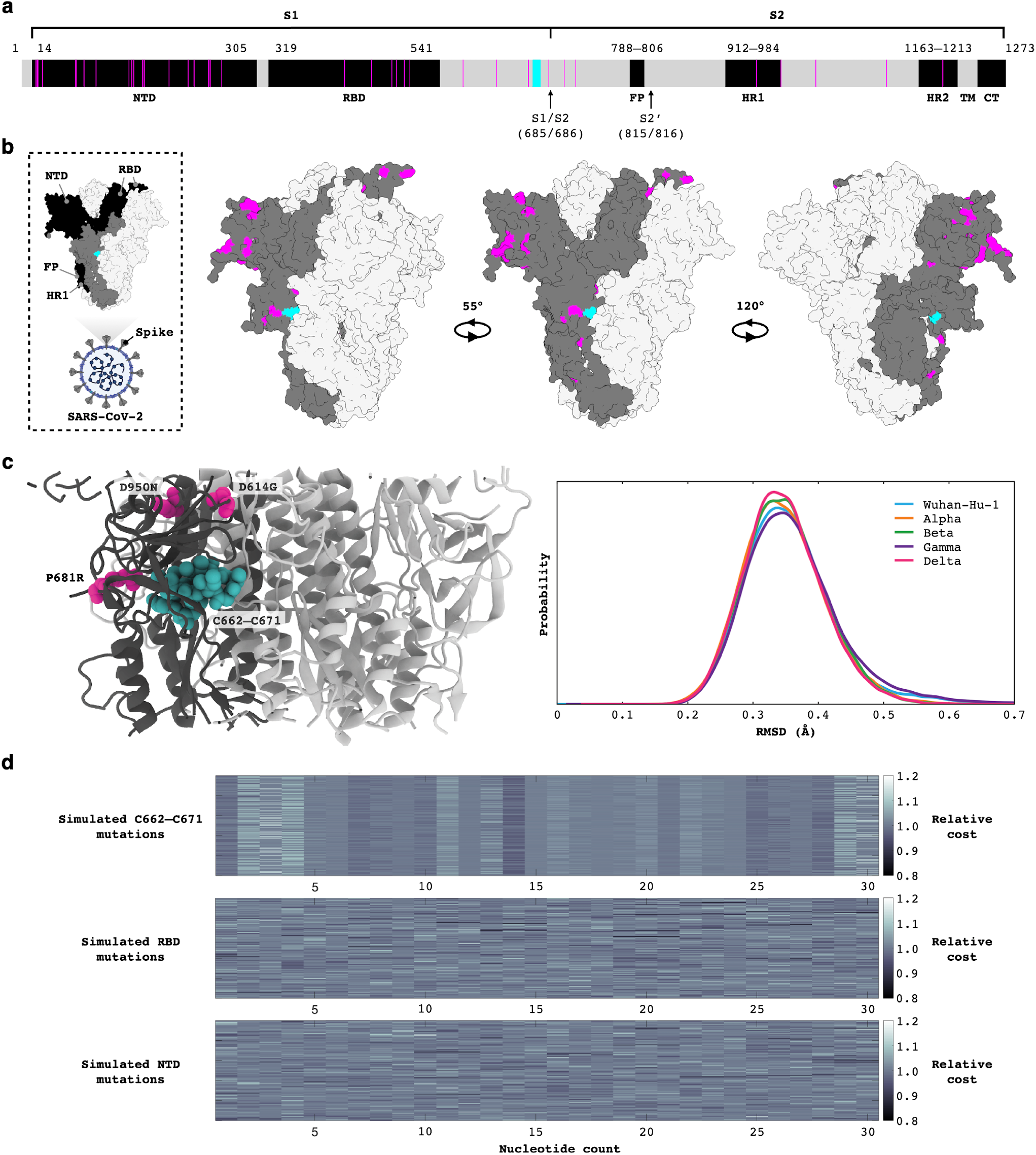
Sequence and structural conservation of the C662–C671 epitope of SARS-CoV-2 S protein across variants. **a**, Domain representation of the missense mutations present throughout the four VOCs (Alpha, Beta, Gamma, and Delta) (magenta) and the unaffected C662–C671 epitope (cyan). The two arrows signify cleavage sites. NTD = N-terminal domain; RBD = receptor-binding domain; FP = fusion peptide; HR = heptapeptide repeat sequence; TM = transmembrane; CT = cytoplasmic tail. **b**, Atomic structural representation of one monomer (dim gray) showing all VOC mutated residues (magenta) and the C662–C671 epitope (cyan). SARS-CoV-2 particle created with BioRender.com. **c**, Each VOC sequence (Alpha, Beta, Gamma, Delta) of the S protein was simulated to assess the impact of mutations on the C662–C671 epitope. The structural model of the Delta VOC that was used for simulations is shown on the left. C662–C671 maintains its conformation found in the Wuhan-Hu-1 strain for all variants. Shown (right) are the probability distributions of the spatial RMSD values (with respect to the Wuhan-Hu-1 strain conformation) of the C662–C671 epitope backbone atoms in each variant, calculated from 500-ns explicit-solvent simulations. **d**, Mutation effects analysis of C662–C671, RBD, and NTD presented as a heatmap of the combined average costs of single nucleotide and amino acid synthesis for the three sequences of interest. For RBD and NTD, 100 sub-sequences of 30-nucleotide length were sampled from the wild-type sequences to generate mutants. The color bar denotes the total cost of nucleotide and amino acid production for mutants normalized to the cost of wild-type production. Each pixel of the heatmap represents a single mutant.

To gain further insight into the structural properties of C662–C671, we performed comparative structural analysis (**Fig. 1b**) and molecular simulations (**Fig. 1c**). The flanking cysteine residues of the C662–C671 epitope interact *via* a disulfide bridge (Cys^1^–Cys^10^) to form a cyclic structure. Prior simulations suggest that such a structural constraint enables the short peptide to retain a conformation with minimal variation.^21^ In addition, all-atom explicit-solvent simulations of the S protein indicate that mutations associated with VOCs do not impact the structural properties of this epitope. For each individual variant, 500-nanosecond (ns) simulations were performed, and the spatial root-mean-square deviation (RMSD) of the C662–C671 epitope was calculated, relative to its configuration in the structure of the original Wuhan-Hu-1 sequence (GenBank accession no. NC_045512.2). For each variant, the probability distribution as a function of RMSD (backbone atoms) was very similar and peaked around 0.35 Å. RMSD values that included the side-chain atoms were also very low and centered around 0.4–0.6 Å (data not shown). These observations strongly suggest that this region remains conformationally stable despite the presence of numerous mutations acquired elsewhere in the protein during the global COVID-19 pandemic. Moreover, the C662– C671 epitope is located on the solvent-exposed surface in both the closed- and open-state of the S protein,^21^ a central structural property for vaccine development, because antibody binding requires access to epitopes. Taken together, these findings support a working hypothesis that the C662–C671 epitope could represent a promising candidate for nearly universal COVID-19 vaccine research and development despite the relatively rapid evolution and selection of SARS-CoV-2 variants. It is also remarkable that the C662–C671 epitope shares 100% identity with the S protein of the previous zoonotic coronavirus (SARS-CoV) that caused the original SARS outbreak in 2003 and partial homology with the corresponding S protein of the Middle East Respiratory Syndrome coronavirus (MERS-CoV) in 2012, indicating the potential role of this epitope for immunization against not only SARS-CoV-2, but also other pathogenic coronaviruses.

Finally, to understand the mechanistic basis of the relatively low mutational rate of the C662–C671 epitope in comparison to the RDB and NTD, we have reasoned that mutations within the two latter protein domains—in addition to providing a functional advantage to the virus—may also be energetically more favorable with respect to the cost of genome replication and translation of mutant sequences. Based on the observation that genome replication and translation combined are the most expensive processes of virus production for the host cells, energy limitation could induce selection pressure and genetic drift on newly incorporated elements in viral genomes.^27^ Thus, the incorporation of “lower-cost” nucleotides or amino acid residues (based on ATP utilization) resulting in energy conservation could be advantageous for SARS-CoV-2.^28^ To experimentally test this possibility, we used the Monte Carlo methodology^29^ to generate mutants of C662–C671, RBD, and NTD and to calculate the cost of synthesis of nucleotides and amino acids relative to the wild-type sequences of these three regions. We have found that the evolutionary costs of production of C662–C671 mutants (both per nucleotide and per residue) are significantly higher than those of RBD and NTD mutants (*p*<0.0001; one-way ANOVA and Tukey’s test) (**Fig. 1d**). Overall, this result indicates that point mutations in the C662–C671 epitope may be evolutionarily costly for the virus compared to mutations in the RBD and NTD, thereby making the former less favorable for natural selection. Such empirical analysis, while admittedly lacking an intimate description of molecular mechanisms involved in virus production, might have captured an essential phenomenon in natural selection of mutations. It also attempts to provide a functional basis for the key observation reported in this study, thereby lending support to the central argument of focusing on either a minimally or non-mutable epitope (towards immunogen universality) of the COVID-19 genome for potentially effective immunization and as a target for inescapable antibodies.

In conclusion, the persistence of the COVID-19 pandemic strongly re-emphasizes the need for adaptable strategies for immunization that enhance efficacy and broaden the spectrum of protection beyond those elicited by the vaccines currently in clinical use and under investigational research and development. We have recently reported that the C662–C671 epitope of SARS-CoV-2 elicits generation of anti-S protein antibodies when incorporated into an aerosol-delivered targeted phage-based vaccine prototype.^20,21^ Here, we refine and integrate structural biology and mathematical tools to present additional experimental evidence that C662–C671 meets all the desirable criteria of an epitope for a potential broad-spectrum, hopefully universal, vaccine against SARS-CoV-2 variants. Moreover, such a versatile approach is highly amenable to simple cloning strategies that would allow display of other exposed and preserved immunogenic epitopes from SARS-CoV-2 or other viral surface glycoproteins. Targeted phage-based vaccines could, therefore, be designed to include admixtures (“cocktails”) of distinct epitopes of the S protein, which may help suppress development of viral resistance or immune evasion.^30^ Finally, targeted phage-based vaccines are affordable to manufacture and robust in cold-free supply chain conditions, which would be particularly impactful in low-income countries with vaccine shortages and distribution challenges. From a broad perspective, this work provides an experimental framework and rational tools for discovery and evaluation of SARS-CoV-2 epitopes with potentially universal features along with the possible generation of inescapable monoclonal antibodies as early therapeutics against COVID-19.

## Materials and Methods

### Atomic Structural Visualization

The atomic structure of S protein was visualized with the University of California, San Francisco (UCSF) Chimera software.^31^ In Fig. 1b, a closed-state structural model of Casalino et al.^32^ was used, which is identical to the cryo-electron microscopy (cryo-EM) model of Walls et al. (PBD ID: 6VXX)^33,34^ except for unresolved loops that have been included (N.B.: residues 1142–1273 removed for visualization purposes).

### SMolecular Dynamics Simulations

To address whether mutations associated with VOCs (Alpha, Beta, Gamma, and Delta) are likely to impact the structural properties of the C662–C671 epitope, all-atom explicit-solvent simulations were performed for the S protein with the associated variant mutant sequences. To focus on the effects of mutations that are near the C662–C671 epitope, simulations were performed for a 5-nm cross section of the S protein trimer (Fig. 1c, left), with position restraints imposed along the cross-sectional boundary. For simulations of the Wuhan-Hu-1 strain sequence, an open-state structural model of Casalino et al.^32^ was used, which is identical to the cryo-EM model of Wrapp et al. (PDB ID: 6VSB)^33,35^ except for unresolved loops that were included. For Alpha, Beta, Gamma, and Delta VOCs, the associated mutations were introduced by using Modeller 10.1.^36^

All-atom explicit-solvent simulations were performed by using GROMACS 2020.3.^37,38^ Each system was solvated by using TIP3P water molecules,^39^ with a 10-Å buffer between the protein and the edge of the box. Na^+^ or Cl^-^ ions were included to neutralize the charge of each system. All simulations employed the AMBER99SB-ILDN force field.^40^ Each system was subsequently subjected to steepest-descent energy minimization, then equilibration under NVT conditions (300 K) for 5 ns and then further equilibration by using the NPT ensemble for 5 ns. During the first round of equilibration, all non-H atoms were position-restrained with harmonic potentials (1000 kJ/nm^2^). All position restraints were removed afterwards for all the atoms outside the cross-sectional boundary and the structure was again energy minimized. A second round of equilibration under NVT conditions at 310 K for 5 ns was then performed and followed by further equilibration with the NPT ensemble for 5 ns. The production simulations were performed for 500 ns (NPT) by using the Parrinello-Rahman barostat^41^ set at 1 bar and the Nosé-Hoover thermostat^42^ at 310 K.

### Calculations and Analysis of Energy Cost for Mutations

Energy can be conserved by using “lower-cost” nucleotides or amino acids (according to ATP utilization). Given that the *de novo* production costs of nucleotides follows the order G>A, C>T, G>C and A>T,^28^ it is not surprising to observe that the SARS-CoV-2 S nucleotide sequence of the Wuhan-Hu-1 strain is AT-rich (nucleotide counts of A: 1,125; C: 723; G: 703; T: 1,271). To quantify the effect of mutations on the change in the cost of nucleotide and amino acid synthesis, we generated mutants from wild-type sequences by randomly selecting a nucleotide from A, T, G, and C to replace the nucleotide at a given position by using the Monte Carlo methodology. Each generated mutant thus differed from the wild type in only one nucleotide position (to mimic point mutations). The probabilities used for random sampling of the nucleotides were based on the nucleotide density of the SARS-CoV-2 S protein (Wuhan-Hu-1 strain, GenBank accession no. NC_045512.2) transcript (*i*.*e*., A: 0.2943; T: 0.3325; G: 0.1839; C: 0.1892) to ensure that the mutants maintained the AT-rich character of its gene sequence. Given that the RBD contains 669 nucleotides and NTD contains 876 nucleotides, the wild-type RBD and NTD were randomly sampled 100 times to select 30-nucleotide sequences to be used for mutant generation (to produce a fair comparison with the 30-nucleotide epitope sequence). For each of the 100 sequences, 10 mutants were generated with respect to each of the 30 nucleotides. As a result, a total of 30,000 mutants each was generated for RBD and NTD. For C662–C671, 1,000 mutants were generated per nucleotide to obtain a total of 30,000 mutants for comparison with RBD and NTD. To quantify the effect of point mutations on cost of nucleotide and amino acid synthesis, for each of the 90,000 mutant nucleotide sequences, the cost (in terms of ATP count) of RNA replication per nucleotide was calculated based on the literature-derived values of ATP required for the synthesis of nucleotides in prokaryotes.^28^ The mutant nucleotide sequences were then translated and the cost of translation per amino acid was calculated based on literature-derived values of amino acid production cost in prokaryotes.^43^ The combined cost of single nucleotide and amino acid production for each mutant was normalized to the combined cost of single nucleotide and amino acid production for the corresponding wild-type sequence. The results were plotted as a heatmap showing the relative cost of each mutant. One-way ANOVA and Tukey’s test were performed to assess the statistical significance of the differences in costs of nucleotide and amino acid production among C662–C671, RBD, and NTD mutants, unless otherwise specified. All analyses were performed in MATLAB R2018a.

## Acknowledgements

We thank Dr. Milka Kostic (Life Science Editors) for professional editorial services. This work was supported in part by core services from Rutgers Cancer Institute of New Jersey (NCI Cancer Center Support Grant P30CA072720); by the RCSB Protein Data Bank, which is jointly funded by the National Science Foundation (NSF) (DBI-1832184), the US Department of Energy (DOE-SC0019749), and the National Cancer Institute, National Institute of Allergy and Infectious Diseases, and National Institute of General Medical Sciences of the National Institutes of Health (NIH) (R01GM133198); by NSF grants (CHE-1614101 and PHY-1522550 to J.N.O. and MCB-1915843 to P.C.W.); and by awards from the Levy-Longenbaugh Donor-Advised Fund (to R.P. and W.A.), and the Welch Foundation (C-1792, to J.N.O.). The work at the Center for Theoretical Biological Physics was also supported by the NSF (Grant PHY-2019745). J.N.O. is a Cancer Prevention Research in Texas Scholar in Cancer Research. The work at Houston Methodist Research Institute was partly supported by the Cockrell Foundation (to P.D.) and the NIH (1R01CA253865 to Z.W. and V.C.; 1R01CA222007 to Z.W. and V.C.; 1R01CA226537 to Z.W., V.C., R.P., and W.A.). We are grateful for the computational resources and support provided by the AMD COVID-19 High Performance Computing (HPC) Fund program, the Northeastern University Discovery cluster, and the Northeastern University Research Computing staff.

## Conflict of Interest

D.I.S., R.P., and W.A. are listed as inventors on a patent application related to this technology (International Patent Application no. PCT/US2020/053758, entitled *Targeted Pulmonary Delivery Compositions and Methods Using Same*). Provisional patent application nos. 63/048, 279, and 63/161,136, entitled *Enhancing Immune Responses Through Targeted Antigen Expression*, have also been filed on the technology and intellectual property reported here. PhageNova Bio has licensed these intellectual properties and C.M., D.I.S., F.H.F.T., T.L.S., S.K.L., R.P., and W.A. may be entitled to standard royalties. S.K.L., R.P., and W.A. are founders and equity stockholders of PhageNova Bio. S.K.L. is a Board Member and R.P. is Chief Scientific Officer and a paid consultant of PhageNova Bio. R.P. and W.A. are founders and equity shareholders of MBrace Therapeutics; R.P. serves as the Chief Scientific Officer and W.A. is a Member of the Scientific Advisory Board at MBrace Therapeutics. These arrangements are managed in accordance with the established institutional conflict-of-interest policies of Rutgers, The State University of New Jersey.

## Author Contributions

C.M., D.I.S., P.D., E.D.-R., F.H.F.T., V.G.C., V.C., P.C.W., S.K.B., J.N.O., R.P., and W.A. designed the research; C.M., D.I.S., P.D., E.D.-R., F.H.F.T., T.L.S., V.G.C., and P.C.W. performed the research; S.K.L. and Z.W. contributed conceptual insights, reagents, or analytical tools; C.M., D.I.S., P.D., E.D.-R., F.H.F.T., T.L.S., V.G.C., Z.W., V.C., P.C.W., S.K.B., J.N.O., R.P., and W.A. analyzed the data; C.M., D.I.S., P.D., F.H.F.T., P.C.W., S.K.B., J.N.O., R.P., and W.A. wrote the initial draft of the manuscript, to which all of the authors contributed edits. J.N.O., R.P., and W.A. supervised the overall project.

